# Age-Dependent Cortical Overconnectivity in Shank3 Mice is Reversed by Anesthesia

**DOI:** 10.1101/2024.08.13.607775

**Authors:** Montagni Elena, Manuel Ambrosone, Alessandra Martello, Lorenzo Curti, Laura Baroncelli, Mannaioni Guido, Francesco Saverio Pavone, Alessio Masi, Anna Letizia Allegra Mascaro

## Abstract

Growing evidence points to brain network dysfunction as a central neurobiological basis for autism spectrum disorders (ASDs). As a result, studies on Functional Connectivity (FC) have become pivotal for understanding the large-scale network alterations associated with ASD. Despite ASD being a neurodevelopmental disorder, and FC being significantly influenced by the brain state, existing FC studies in mouse models predominantly focus on adult subjects under anesthesia. The differential impact of anesthesia and age on cortical functional networks in ASD subjects remains unexplored. To fill this gap, we conducted a longitudinal evaluation of FC across three brain states and three ages in the Shank3b mouse model of autism. We utilized wide-field calcium imaging to monitor cortical activity in Shank3b^+/-^ and Shank3b^+/+^ mice from late development (P45) through adulthood (P90), and isoflurane anesthesia to manipulate the brain state. Our findings reveal that network hyperconnectivity, emerging from the barrel-field cortices during the juvenile stage, progressively expands to encompass the entire dorsal cortex in adult Shank3b^+/-^ mice. Notably, the severity of FC imbalance is highly dependent on the brain state: network alterations are more pronounced in the awake state and shift towards hypoconnectivity under anesthesia. These results underscore the crucial role of anesthesia in detecting autism-related FC alterations and identify a significant network of early cortical dysfunction associated with autism. This network represents a potential target for non-invasive translational treatments.

## INTRODUCTION

Autism spectrum disorders (ASDs) are a group of neurodevelopmental conditions characterized by deficits in social communication, interaction, and repetitive behaviors. Emerging evidence suggests that brain network dysfunction are central to the neurobiological basis of ASDs. Thus, functional connectivity (FC) studies are pivotal for exploring the interactions between brain regions and elucidating autism-related large-scale network dynamics.

Research has identified a range of hyper- and hypo-connected networks as primary alteration in adult mouse models of autism (Balasco et al. 2022; Hull et al. 2018; Zerbi et al. 2022; Nakai et al. 2023). However, behavioral deficits and brain circuits anomalies are already evident in early and late postnatal phases (Peixoto et al. 2019), which are crucial periods for effective intervention (Chung, Shin, and Kim 2022; Qin et al. 2018). While functional imaging during developmental stages has shown predictive value for autism diagnosis in humans (Emerson et al. 2017), similar longitudinal studies in mice are lacking, particularly those that track FC from development through adulthood (Peixoto et al. 2019).

Furthermore, FC is influenced by brain state (Finn et al. 2017; Stitt et al. 2017) which in turn is frequently constrained by the imaging method used (Paasonen et al. 2018). Functional Magnetic Resonance Imaging (fMRI) is the preferred technique for FC studies in mice due to its strong translational potential to humans. However, fMRI is typically conducted under anesthesia to prevent motion artifacts (Jonckers et al. 2015; Lambers et al. 2023). Anesthesia can preserve some functional networks while suppressing or altering others (Paasonen et al. 2018), making it challenging to validate FC results across different brain states. Although recent advances have enabled fMRI in awake head-fixed autistic mice (Tsurugizawa et al. 2020; Mandino et al. 2024), mesoscale Ca^2+^ imaging offers a valuable alternative for investigating the cortical network across different brain states within the same subjects (Montagni et al. 2024; Scaglione et al. 2024). Despite its limitations to near-surface recordings (Nietz et al. 2022), this technique combines wide-field fluorescent microscopy (WFFM) with genetically encoded calcium indicators to directly measure neuronal activity (Resta et al. 2022; Montagni et al. 2018).

WFFM has recently been used to explore altered FC patterns during voluntary movements in the Shank3 mouse model of autism (Nakai et al. 2023). SHANK3 is a postsynaptic scaffolding protein of excitatory synapses, whose reduced expression leads to impaired synaptic transmission and plasticity (Peixoto et al. 2019; Chung, Shin, and Kim 2022). It is known to cause Phelan-McDermid syndrome (PMS) and idiopathic autism (Costales and Kolevzon 2015; De Rubeis et al. 2018; Monteiro and Feng 2017). Adult Shank3b mutant mice exhibit anxiety-like behavior, excessive self-injurious grooming, and altered whisker-dependent discrimination (Balasco et al. 2022; Peça et al. 2011; Orefice et al. 2019), suggesting significant cognitive and sensory dysfunctions.

In this study, we investigated the brain state-dependence of cortical network dynamics using mesoscopic Ca^2+^ imaging of excitatory neurons expressing GCaMP7f in Shank3b^+/+^ and Shank3b^+/-^ mice. We longitudinally assessed cortical network alterations at three developmental stages, beginning at postnatal day 45 (P45) and extending to adulthood (P90). Our findings indicate that hyper-connectivity in the barrel cortices contributes significantly to the emergence of aberrant FC networks from juvenile stages, and a progressive strengthening and expansion of this overconnected network with age. Importantly, these network dysfunctions are not sustained under deep anesthesia. Theseresultsreveal a critical network of cortical dysfunction associated with autism, which may serve as a target for noninvasive translational therapies.

## RESULTS

In this study, we first examined behavioral alterations in Shank3b ^+/-^ mice at a juvenile stage (P20). We then assessed FC impairments at multiple developmental stages, from late adolescence (P45) to adulthood (P90), using WFFM. To understand how cortical network alterations were influenced by brain state, we compared FC across different conditions: wakefulness, light anesthesia, and deep anesthesia. This approach allowed us to evaluate the impact of varying brain states on the altered cortical network.

### Adolescent Shank3b^+/-^ Mice Exhibit Anxiety and Altered Grooming Behavior

Behavioral assessments of Shank3b mutant mice at various ages starting from late postnatal development (P45 to 6 months) have previously identified cognitive dysfunctions (Balasco et al. 2022; Orefice et al. 2019). To confirm behavioral anomalies present in our mouse mutant model, we investigated Shank3b ^+/-^ mice at postnatal day 20 (P20, fig.1a). Generalized anxiety and locomotor activity were evaluated by using an open field arena. Although Shank3b^+/-^ mice traveled similar distances compared to their Shank3b^+/+^ littermates (fig.1b), they spent less time in the center of the arena (fig.1c), indicating heightened anxiety-like behavior without observable motor deficits. In line with previous study (Orefice et al. 2019; Liu et al. 2024), the total grooming time did not significantly differ between genotypes (fig.1d). However, Shank3b^+/-^ showed a reduced frequency of grooming episodes (fig.1e) and a notable tendency to engage in longer grooming bouts (fig.1f). Thesefindings suggest that while overall grooming activity remains unchanged, Shank3b^+/-^ mice exhibit difficulties in interrupting repetitive behaviors, highlighting a specific grooming dysfunction.

**Figure 1.**
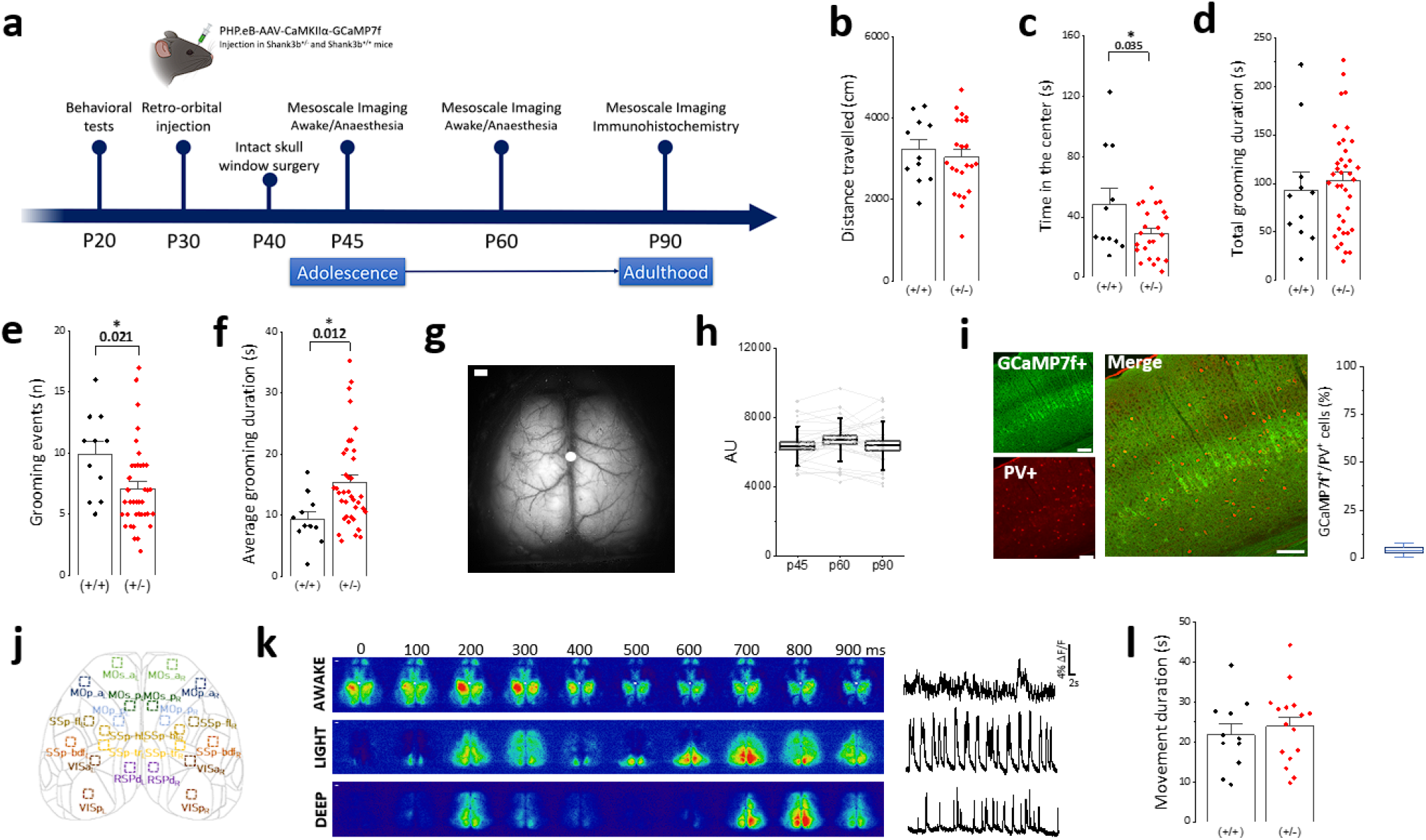
Experimental design for cortical mesoscopic calcium imaging, immunohistochemical analysis and behavioral characterization of Shank3b+/-mice. **(a)** Experimental timeline, with behavioral test at P20 followed by retro-orbital injection of AAV-PHP.eB GCaMP7f at P30, optical window implantation at P40 and longitudinal imaging timepoints at P45, P60 and P90. At the end of in vivo experiments, immunohistochemical analysis was performed. **(b, c, d, e, f)** Quantification of open field performance of Shank3B+/+ and Shank3B+/-mice, **(b)** Total distance travelled (Shank3b+/+, 3220 ± 240 cm n=11; Shank3b+/-3034 ± 188 cm n=23, two-sample t-test), **(c)** Time spent in the center of the arena (Shank3b+/+ 48 ± 10 s n=11; Shank3b +/-29 ± 3 s n=11, *p<0.05, two-sample t-test), **(d)** quantification of the total amount of self-grooming grooming (93 ± 18 s n=11 Shank3b+/+, 103 ± 8 s n=38 Shank3b +/-, two-sample t-test), **(e)** quantification of the number of self-grooming events (n) performed (9.9 ± 1.0 n n=11 Shank3b+/+, 7.1 ± 0.5 n=38 Shank3b +/-, *p<0.05, two-sample t-test). **(f)** Averaged duration of a single self-grooming event (9 ± 1 s n=11 Shank3b+/+, 15 ± 1 s n=38 Shank3b +/-, *p<0.05, two-sample t-test). **(g)** Representative image of the field of view. White dots represent bregma, scalebar: 1mm. **(h)** In vivo quantification of GCaMP7f expression across timepoints (6326 ± 223 AU - P45, 6679 ± 245 AU - P60, 6354 ± 277 AU - P90, n = 27, one-way repeated measure ANOVA). **(i)** Left, representative immunohistochemistry images showing the neuronal expression of GCaMP7f (green) and PV (red) in the whole cortical layers, scalebar: 100 µm. Right, Quantification of the colocalization ratio GCaMP7f+/PV+ (4.0 ± 1.5 %, n=6 FOV in 1 mouse). **(j)** Cortical parcellation map with 10×10 ROI based on Allen Mouse Brain Atlas (see Methods). **(k)** Left, representative image sequences showing cortical activity in three different brain states (awake, light anesthesia and deep anesthesia). White dots represent bregma, scale bar: 1 mm. Right, representative single -trail time series showing averaged cortical activity in the same brain states. **(l)** Quantification of forepaw movement during imaging session at P45. Shank3B+/+ (black) and Shank3B+/-(red) spent the same time doing movement (21.8 ± 2.6 s, n=11 Shank3b+/+; 23.9 ± 2.3s, n=16 Shank3b+/-, two-sample t-test). Data represent mean ± SEM, each dot represents one animal. All plots report the mean values ± SEM, each dot represents one animal.

### AAV-PHP.eB Delivered at Postnatal Day 30 Induces Long-term Stable Transgene Expression in the Cortex

To investigate the cortical patterns underlying functional impairments observed in adolescent Shank3b^+/-^ mice, we employed retro-orbital (RO) injections of a recently developed viral vector able that effectively crosses the blood-brain barrier (BBB) (Michelson, Vanni, and Murphy 2019; Chan et al. 2017). The viral vector, AAV-PHP.eB-CaMKIIα-GCaMP7f, enables widespread transduction of the calcium indicator throughout the cortex, allowing for comprehensive detection of excitatory neuronal activity across the dorsal cortical mantle (fig.1g). We assessed the long-term stability of GCaMP7f expression by injecting AAV-PHP.eB-CaMKIIα-GCaMP7f at P30 and measuring spatial fluorescence intensity profiles at three time points, i.e., P45, P60 and P90. GCaMP7f expression remained stable through P90 (fig.1h). To verify the specificity of CaMKII*α* promoter for excitatory neurons, we used an antibody against parvalbumin (PV) assessing the density of PV interneurons in post-mortem brain sections from P90 mice (fig.1i). Our results confirmed that the majority of PV-interneurons were not transfected, validating the promoter selectivity for targeting excitatory neurons (fig.1i).

This approach allowed us to capture cortical activity from 22 regions of interest (ROIs) across the entire dorsal cortex (fig.1j) during different brain states: wakefulness, light anesthesia and deep anesthesia (fig.1k). During the awake state, mice were head-restrained and free to move their limbs without specific tasks. The average time spent in movement episodes during a 3-minute awake imaging session was comparable between Shank3b^+/-^ and Shank3b^+/+^ mice (fig.1l). Anesthesia was modulated by adjusting isoflurane concentration between 1.3% to 1.7% in accordance with previous studies (Montagni et al. 2024). Light anesthesia resulted in longer Up-state durations (Supp. Fig.1b-d) and shorter Down-state durations (Supp Fig.1a-c) compared to deep anesthesia in both genotypes. Up- and Down-state durations remained consistent across different time points (Supp Fig.1). These data demonstrate that systemic administration of AAV-PHP.eB effectively induces selective and long-term stable expression of calcium indicator at the cortical level, enabling longitudinal mesoscale Ca ^2+^ imaging in vivo in a mouse model of autism.

### Awake State Hyperconnectivity of Barrel Field and Visual Cortices in Adolescent Shank3b^+/-^ Mice

We first applied our approach to investigate cortical FC alterations in P45 Shank3b ^+/-^ mice and their dependency on brain state. By characterizing FC at P45, we aimed to determine if early FC of excitatory neurons exhibited region-specific changes in Shank3b^+/-^ mice compared to their non-mutant littermates, and whether these abnormalities were preserved across different brain states. FC matrices were obtained by calculating the Pearson’s correlation coefficient (Fisher’s z-transformed) between time traces of paired cortical regions after hemodynamic correction (Fig.2a-c). We then computed differences between Shank3b^+/+^ and Shank3b^+/-^ mice within each brain states, obtaining a matrix of differences (Fig.2d-f), and tested for significant changes in correlations using the Network-Based Statistic (NBS, Fig.2g) (Nakai et al. 2023).

**Figure 2.**
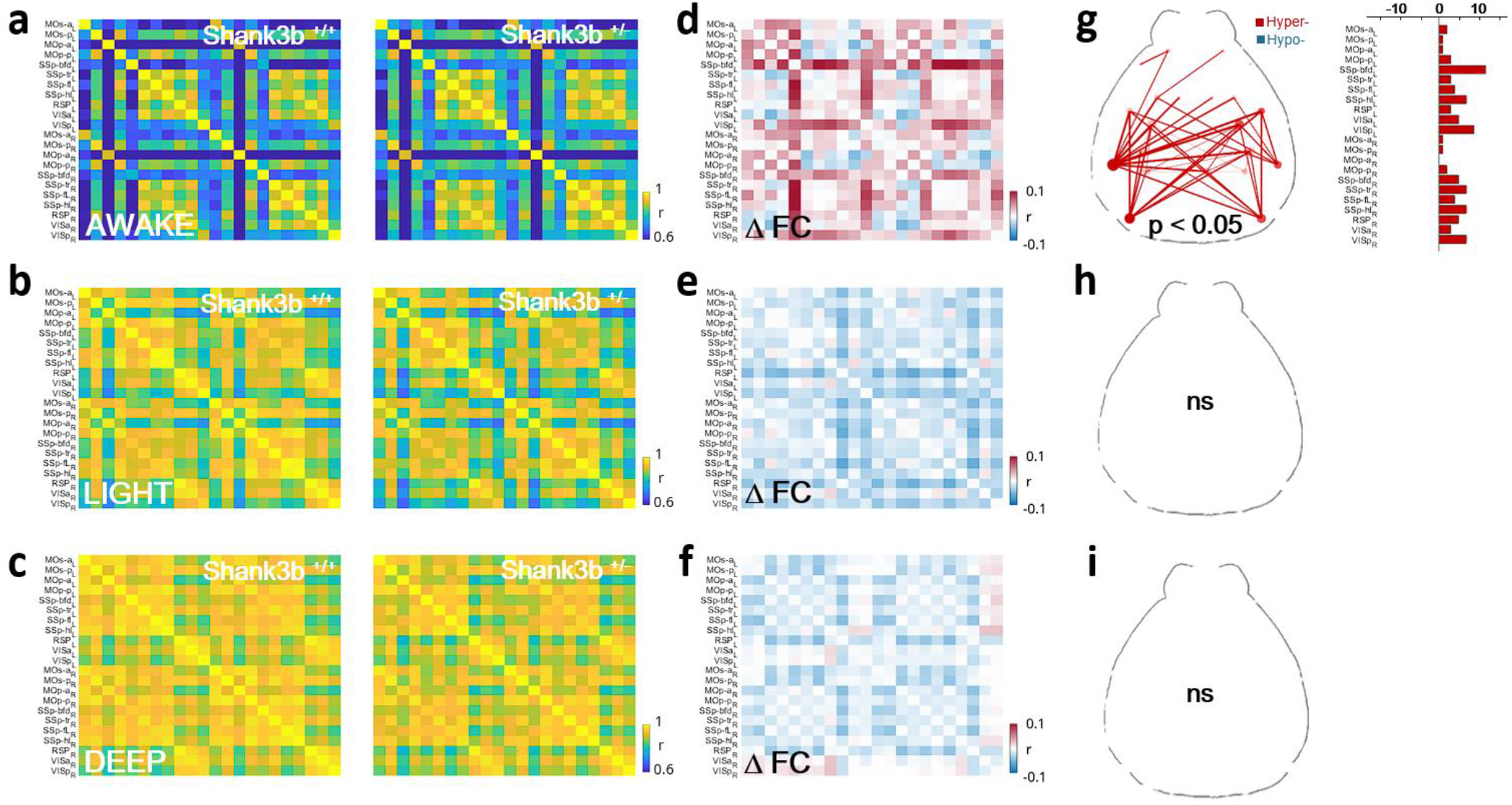
Awake state hyperconnectivity of barrel field and visual cortices in adolescent Shank3b+/-mice (P45) **(a** and **b** and **c)** Pairwise Pearson’s correlation coefficients of cortical activity in a three minutes window were visualized as averaged correlation matrices for genotype (Shank3b+/+ left, Shank3b+/-right) in the awake state **(a)**, light anesthesia **(b)** and deep anesthesia **(c). (d** and **e** and **f)** Matrix of difference, produced by subtracting the average FC of Shan3b+/+ from that of Shank3b+/ - mice in the awake state **(d)**, light anesthesia **(e)** and deep anesthesia **(f).** Red and blue squares indicate Shank3b+/- hyper- or hypo-connectivity respectively. **(g** and **h** and **i)** Network diagrams (left) of statistically significant FC alterations in the awake state **(g)**, light anesthesia **(h)** and deep anesthesia **(i)**. The bar plot (right) indicates the number of significant FC alterations for each cortical area. (n = 11 Shank3b+/+ and 17 Shank3b+/ -).

Consistent with previous studies (Scaglione et al. 2024), resting-state (rsFC) increased with higher levels of anesthesia (Fig.2a-c). Although both genotypes exhibited this trend, genotype-specific effect were observed. A robust and symmetric hyperconnectivity was evident in the awakestate of Shank3b^+/-^ mice compared to Shank3b^+/+^ mice (Fig.2d). This hyper-connected network primarily involved the barrel field (SSp-bfd) and visual cortices (VISp, Fig.1g), suggesting that the altered sensitivity to peripheral stimuli observed in Shank3b mutant mice (Balasco et al. 2022) may arise from early network impairment. Under light (Fig.2d) and deep anesthesia (Fig.2f) Shank3b^+/-^ FC was instead similar to control mice, showing a non-significant trend toward global hypo-connectivity (Fig.2e-f).

The substantial dependence of FC alterations on the brain state is indicated by the formation of a hyper-connected network in the waking state of Shank3b^+/-^ mice, which fades under anesthesia. These results underscore how crucial is to monitor the brain state when comparing FC between mutant mice and their wild-type littermates.

### Hyperconnectivity Becomes More Prominent and Spreads to Motor Regions at P60

To evaluate how the cortical network evolves with age, mice we monitored at P60. Interestingly, the hyper-connected network involving the SSp-bfd strengthened in the awake state of Shank3b^+/-^ mice at P60 (Fig.3d-g). At this stage there was also an emergence of impairment in the anterior primary motor cortices (MOp -a) in Shank3b^+/-^ mice (Fig.3g). In contrast to the results observed at P45, hyperconnectivity also emerged under light anesthesia at P60 (Fig.3e). These significant alterations affected the entire cortical network, except for the retrosplenial cortices (RSP, Fig.3f). Under deep anesthesia, there was a widespread but mild reduction in FC, with no significant alterations (Fig.3g-h). Collectively, these results highlight the persistence and strengthening of a densely hyper-correlated subnetwork involving the barrel field cortex of both hemispheres over time in the awake state. Additionally, the spread of hyperconnectivity to motor regions at P60 indicates a progression of network dysfunction as the mice age.

**Figure 3.**
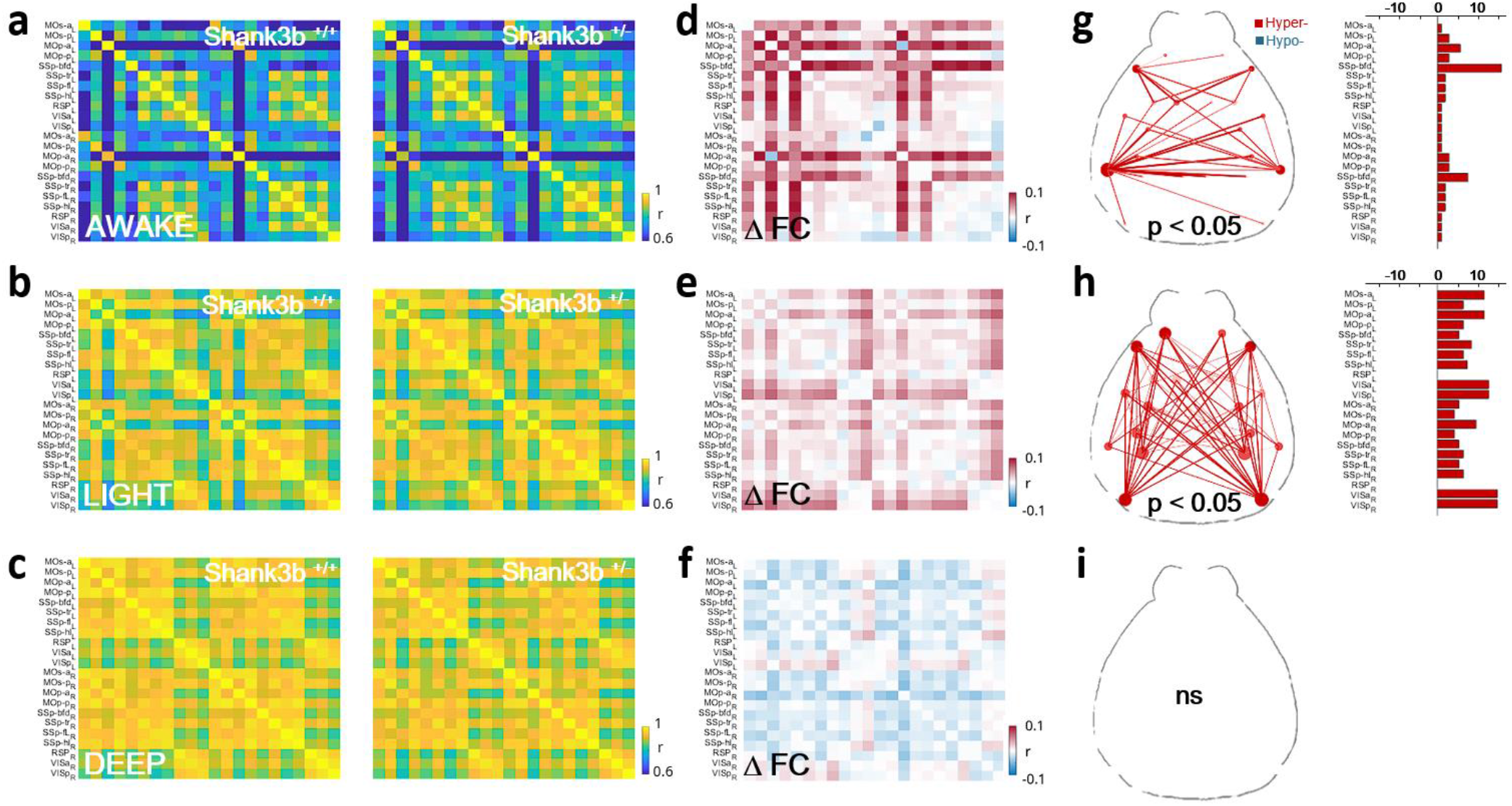
Hyperconnectivity becomes more prominent and spreads to motor regions at P60. **(a** and **b** and **c)** Pairwise Pearson’s correlation coefficients of cortical activity in a three-minute window were visualized as averaged correlation matrices for genotype (Shank3b+/+ left, Shank3b+/-right) in the awake state **(a)**, light anesthesia **(b)** and deep anesthesia **(c). (d** and **e** and **f)** Matrix of difference, produced by subtracting the average FC of Shan3b+/+ from that of Shank3b+/ - mice in the awake state **(d)**, light anesthesia **(e)** and deep anesthesia **(f).** Red and blue squares indicate Shank3b+/- hyper- or hypo-connectivity respectively. **(g** and **h** and **i)** Network diagrams (left) of statistically significant FC alterations in the awake state **(g)**, light anesthesia **(h)** and deep anesthesia **(i)**. The bar plot (right) indicates the number of significant FC alterations for each cortical area. (n = 13 Shank3b+/+ and 17 Shank3b+/ -).

### Alterations in Functional Connectivity Persist at P90

We monitored alterations in FC from adolescence to adulthood to assess their persistence. In the awake state at P90, a dense and widespread hyper-correlated cortical network emerged (Fig.4d), indicating that differences in FC between the two genotypes increased over time. Hyperconnectivity also persisted in lightly anesthetized P90 mice (Fig.4e). However, under deep anesthesia, Shank3b^+/-^ mice exhibited a non-significant trend towards hypo-connectivity compared to Shank3b^+/+^ mice (Fig.4f). Taken together, these results suggest a robust dependence of the altered network on the brain state, enduring from P45 to P90. The progression of hyperconnectivity in awake and lightly anesthetized states underscores the persistent and evolving natureof network dysfunction in Shank3b^+/-^ mice.

**Figure 4.**
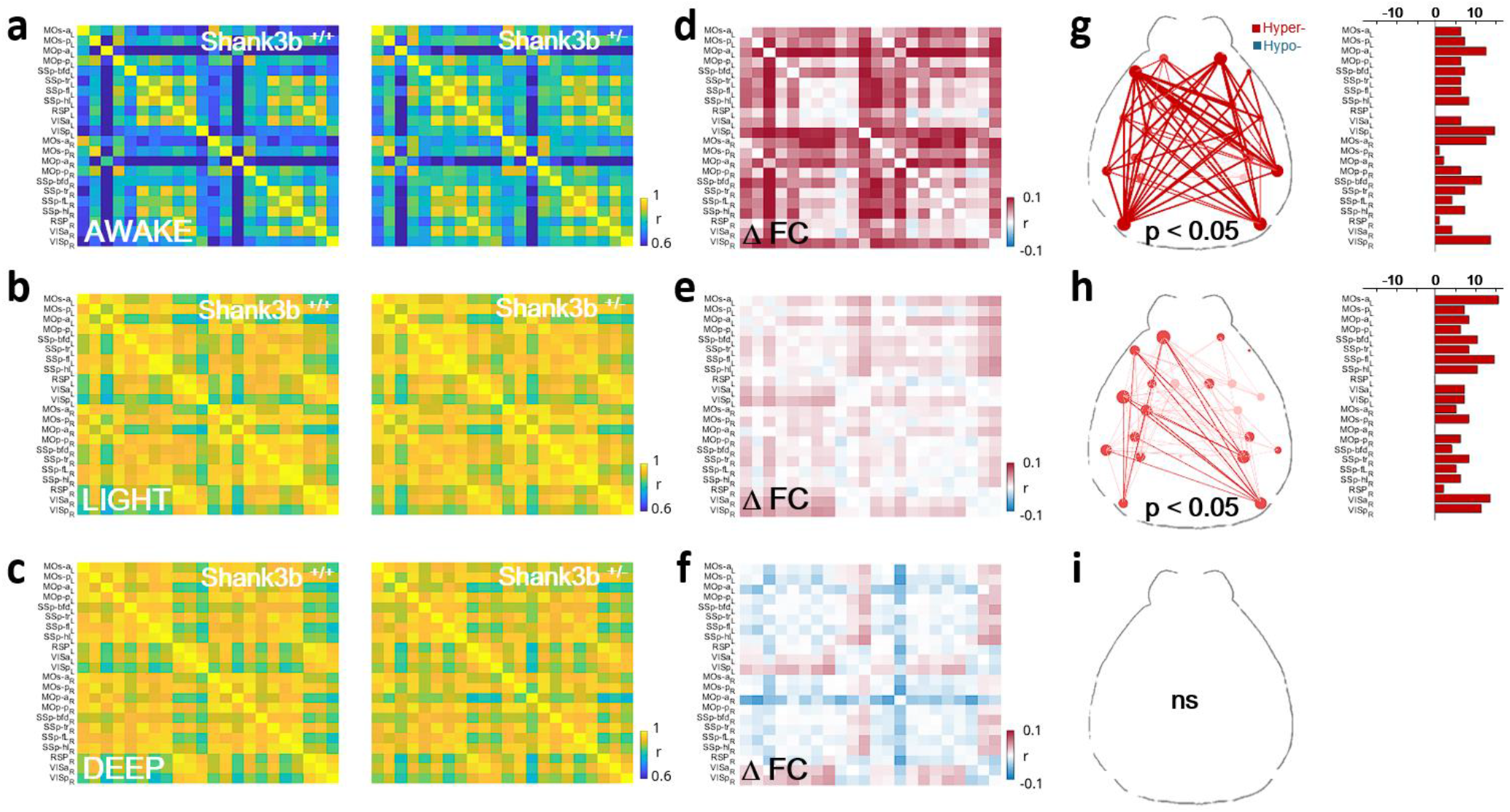
Alterations in functional connectivity persist at P90. **(a** and **b** and **c)** Pairwise Pearson’s correlation coefficients of cortical activity in a three minutes window were visualized as averaged correlation matrices for genotype (Shank3b+/+ left, Shank3b+/ - right) in the awake state **(a)**, light anesthesia **(b)** and deep anesthesia **(c). (d** and **e** and **f)** Matrix of difference, produced by subtracting the average FC of Shan3b+/+ from that of Shank3b+/- mice in the awake state **(d)**, light anesthesia **(e)** and deep anesthesia **(f).** Red and blue squares indicate Shank3b+/- hyper- or hypo-connectivity respectively. **(g** and **h** and **i)** Network diagrams (left) of statistically significant FC alterations in the awake state **(g)**, light anesthesia **(h)** and deep anesthesia **(i)**. The bar plot (right) indicates the number of significant FC alterations for each cortical area. (n = 13 Shank3b+/+ and 14 Shank3b+/-).

### Network Alterations are Strongest in the Awake State

Given that FC alterations intensified and spread across the entire cortex between P45 and P90 in both the awake state and under light anesthesia, we compared the temporal evolution of the average global FC between genotypes in each brain state separately.

Interestingly, Shank3b^+/+^ global FC remained stable over time in all three brain states investigated (fig. 5). In contrast, Shank3b^+/-^ global FC significantly increased from P45 to P90 in both the awake state and under light anesthesia (fig. 5a). Under deep anesthesia, global FC trends in Shank3b ^+/-^ were similar to those in Shank3b^+/+^ mice, with no significant differences (fig. 5c). It is important to note that FC changes under anesthesia are not influenced by the brain state, as the Up- and Down-state durations were consistent across time and genotypes (Supp Fig1 a-b). Further analysis revealed that FC alterations between Shank3b^+/+^ and Shank3b^+/-^ were more pronounced in the awake state and intensified over time (Fig. 5d-f). This demonstrates a robust hyper-connectivity of the cortical network in awake Shank3b^+/-^ mice starting from a juvenile age. Moreover, these results highlighted that the severity of connectivity imbalance is highly dependent on the brain state: network alterations are more pronounced in the awake state and tend towards hypoconnectivity under anesthesia (Fig. 5d-f).

**Figure 5.**
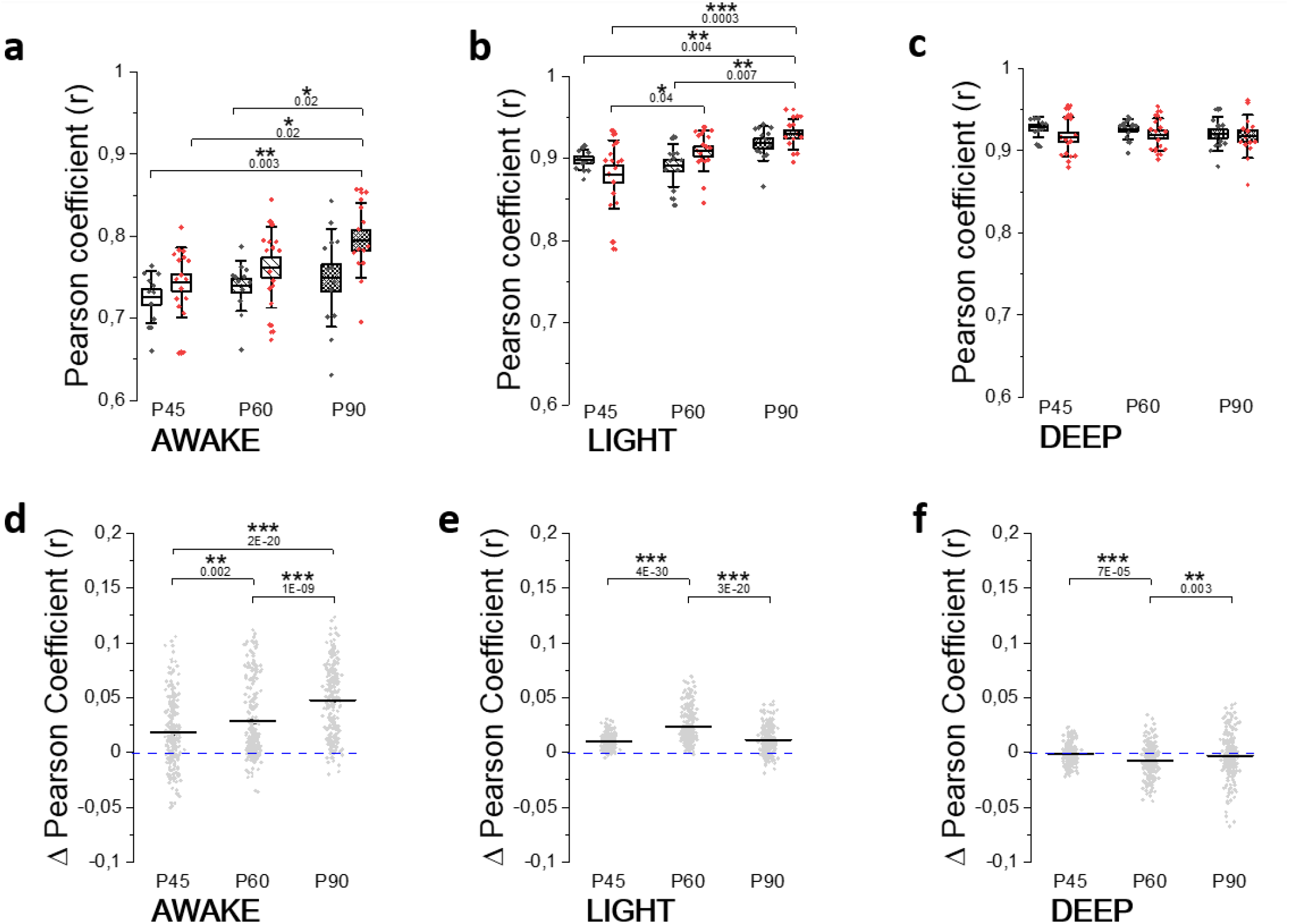
Network alterations are stronger in the awake state and increase over time. **(a-c)** Box plots of global functional connectivity across time for both the genotypes (Shank3b+/+ black, Shank3b+/-red) in the awake state **(a)** light anesthesia **(b)** and deep anesthesia **(c)** (* p < 0.05; ** p < 0.01; *** p < 0.001, two-way ANOVA). **(d-f)** Longitudinal monitoring of difference in person coefficient between Shank3b(+/-) and Shank3b(−/-) in awake **(d)**, light **(e)** and deep anesthesia **(f)**.

**Figure 6:**
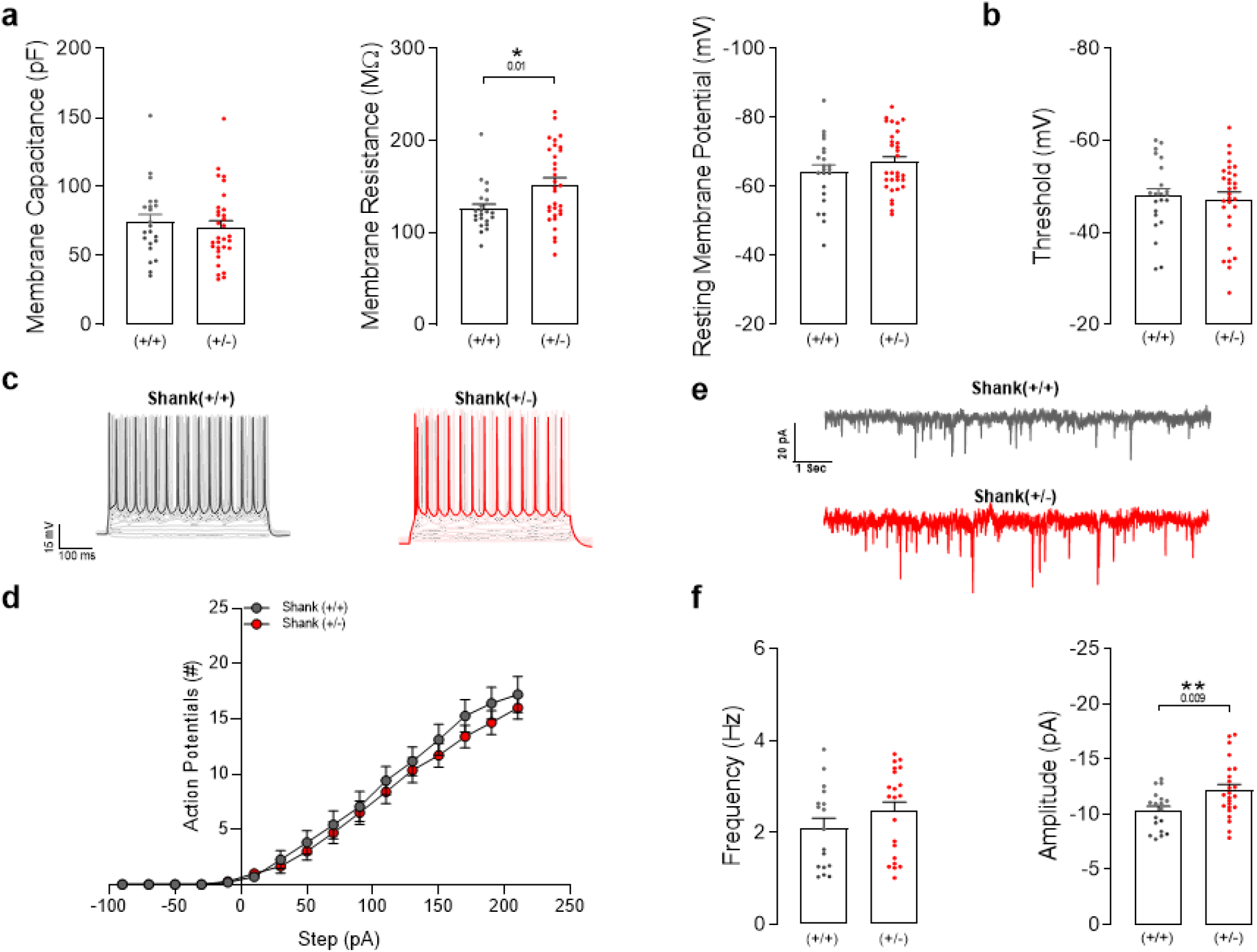
Characterization of layer V pyramidal neurons of SSp-bfd. **(a)** Passive properties such as membrane capacitance (left, Shank3b+/+, 74.28 ± 5.65 pF, N=7 mice, n=22 cells; Shank3b+/-, 70.53 ± 4.757 pF, N=8 mice, n=31 cells), membrane resistance (middle, Shank3b+/+, 125.6 ± 5.32 mΩ, N=7 mice, n=22 cells; Shank3b+/-, 152.3 ± 7.45 mΩ, N=8 mice, n=31 cells), and resting membrane potential (right, Shank3b+/+, -64.2 ± 2.069 mV, N=7 mice, n=22 cells; Shank3b+/-, -67.06 ± 1.58 mV, N=8 mice, n=31 cells) in Shank3b+/+ (black) and Shank3b+/-mice (red). **(b)** Action potential threshold (Shank3b+/+, -47.89 ± 1.67 mV, N=7 mice, n=21 cells; Shank3b +/-, -47.24 ± 1.57 mV, N=8 mice, n=31 cells). (**c)** Representative current clamp traces obtained by 500 ms steps of increasing depolarizing current ranging from -90 to +210 in 20pA increments. Bold traces show the action potential firing at 210 pA current input in Shank3b+/+ (grey) and Shank3b+/-(red). (**d)** Current-AP curves obtained by plotting the mean number of AP evoked against current input amplitude (Shank3b+/+, N=7 mice, n=20 cells; Shank3b +/-, N=8 mice, n=26 cells). (**e)** Representative whole-cell recordings of sEPSCs from Shank3b+/+ (grey) and Shank3b+/-(red) mice. **(f)** Bars graph of absolute (dots) and average (bars) measure of sEPSCs frequency (left, Shank3b+/+, 2.095 ± 0.218 Hz, N=7 mice, n=17 cells; Shank3b +/-, 2.460 ± 0.938 Hz, N=8 mice, n=21 cells) and amplitude (right, Shank3b+/+, -10.32 ± 0.4184 pA, N=7 mice, n=18 cells; Shank3b +/-, -11.88 ± 0.3829 pA, N=8 mice, n=24 cells) in both genotypes.

### Elevated Response to Spontaneous Excitatory Inputs in the SSp-bfd of Adult Shank3b+/-Mice

Given the alterations in large-scale excitatory FC, we targeted the underlying synaptic dysfunctions in SSp-bfd neurons at P90. Whole-cell recordings were conducted on *ex-vivo* brain slices from Shank3b+/+ and Shank3b+/-mice, specifically targeting layer V pyramidal neurons of the SSp-bfd. Firstly, we evaluated the intrinsic electrophysiological properties and detected no changes in the membrane capacitance and resting membrane potential. However, a significant increase in membrane resistance was identified in Shank3b^+/-^ mice (Fig.5a).

Subsequently, we examined theintrinsicexcitability of layer V pyramidal neurons by measuring the action potential threshold and the number of action potentials evoked by depolarizing steps of increasing amplitude (Fig.5d). As shown in Fig.5b-c, we found no differences in these measures, suggesting that the genotype does not influence somatic responses to current stimuli. Recordings of spontaneous excitatory postsynaptic currents (sEPSCs) revealed no difference in the frequency of sEPSCs, while an increased sEPSCs amplitude was found in Shank3b^+/-^ compared to wild-type littermates (Fig.5f).

This indicates that while the rate of spontaneous synaptic events remains unchanged, the postsynaptic response to these events is significantly enhanced in Shank3b^+/-^ mice, suggesting a heightened sensitivity to excitatory inputs in the SSp-bfd neurons. These findings provide a potential synaptic mechanism underlying the observed hyperconnectivity in the cortical network of Shank3b^+/-^ mice, emphasizing the role of enhanced synaptic strength in the altered FC associated with this model of autism.

## DISCUSSION

This study aimed to address the gap in autism-related FC research, which predominantly focuses on adult stages. Autism is a neurodevelopmental disorder, yet there is a scarcity of studies examining brain connectivity during development in rodent models. Additionally, how brain states differentially affect cortical functional networks in autistic subjects remains largely unexplored.

Since SHANK3 haploinsufficiency is responsible for the neurobehavioral phenotype in individuals with Phelan-McDermid Syndrome (PMS), we focused on Shank3 heterozygous mice generated by crossing wild -type mice with heterozygotes to be most relevant to the clinical syndrome. Previous research has mainly focused on Shank3b knockout mice (Shank3b^-/-^) due to their more pronounced behavioral defects. However, heterozygous SHANK3 mutations or loss are considered a major cause of PMS in humans (Costales and Kolevzon 2015). Thus, Shank3b^+/-^ mice offer a more relevant model for studying cortical FC alterations.

We used mesoscale cortical imaging to study excitatory neuron activity in Shank3b^+/-^ mice, tracking changes from late adolescence to adulthood. We utilized retro-orbital injection of AAV-PHP.eB-GCaMP7f at P30, achieving effective and stable transduction of excitatory neurons across the dorsal cortex with a strong tropism for layer V (L5) pyramidal neurons.This method provides a reliable tool for monitoring cortical activity across ages (Michelson, Vanni, and Murphy 2019; Mathiesen et al. 2020) and avoids the high costs and time associated with cross-breeding genetically modified mice with GECI-expressing strains (Grødem et al. 2023).

Behavioral assessments revealed generalized anxiety and repetitive behaviors in young Shank3b^+/-^ mice (P20), consistent with previous findings of neophobia and over-grooming in Shank3b^+/-^ mice at later stages (Orefice et al. 2019; Jaramillo et al. 2017; Peça et al. 2011). Moreover, we found that early alterations in the cortical somatosensory network were evident by P45, persisting until adulthood (P90). These results support the hypothesis that somatosensory connections are already well-established before adolescence (P15) (Rahn et al.2021) and that an early excitation/inhibition (E/I) imbalance and myelin defects during the early stage of development – before P21 - may be an essential mechanism leading to alterations in behavior and FC (Malara et al. 2022; Fotiadis et al. 2023; Rasero et al. 2023). Consistently, local network excitability was impaired at P90, with an increased amplitude of sEPSCs in L5 barrel cortex pyramidal neurons from Shank3^+/-^ mice. Of note, the significant increase in membraneresistanceassociated with the Shank3^+/-^ genotypedose not translate into a change in intrinsic somatic excitability.

Altered connectivity in the motor cortices emerged later (P60), likely reflecting late maturation processes (Rahn et al. 2021). Indeed, anterior portions of the brain, including prefrontal cortices, undergo late maturation: motor, visual and retrosplenial cortices connections significantly changed between P22 and P60 in healthy mice (Rahn et al. 2021; Mallya et al. 2019). These findings align with reports of hyper-connected cortical patterns involving motor areas in other autism mouse models during wakefulness (Nakai et al. 2023).

The progressive alterations in FC from P45 to P90 may result from a gradual deterioration of circuit function with age, which is consistent with the high rate of regression reported in PMS patients. Neurodevelopmental regression is a key feature of PMS (Dille et al. 2023), with affected individuals showing a distinctive pattern of regression through several stages across the lifespan. Notably, regression in PMS primarily affects motor and self-help skills (Reierson et al. 2017), which corresponds with our findings that indicate a later involvement of motor cortices in FC alterations. However, it is important to note that rs-fMRI studies have shown hypo-connectivity within the somatosensory-hippocampal network and prefrontal regions in Shank3b^-/-^ mice under anesthesia (Balasco et al. 2022; Pagani et al. 2019).

Consolidating our previous work on the brain state dependence of the FC in healthy subjects (Montagni et al. 2024), this study demonstrates how brain states significantly impact network dynamics in autistic mice. Across all investigated time points, there is a discernible shift from hyper-to hypo-connectivity in Shank3b^+/-^ mice compared to Shank3b^+/+^ mice when transitioning from wakefulness to anesthesia. This finding reconciles the seemingly conflicting results from different studies, highlighting the crucial role of brain state in modulating cortical functional connectivity in this autism model.

Based on these findings of abnormally higher connectivity in awake Shank3b^+/-^ mice, we speculate that the cortical network of the mutants does not reach full maturation, possibly due to developmental miswiring (Di Martino et al., Neuron 2014). Previous studies have shown that the dynamic process of over-connectivity followed by pruning, which occurs at neuronal level, also operates at the systems level, fostering the reshaping and balancing of connectivity in the developing brain (Supekar et al., Plos Biol 2009). Accordingly, brain networks in children present lower levels of hierarchical organization and modularity (Fair et al., Plos Comp Biol 2009). In the incompletely mature brains of Shank3b^+/-^ mice, this could result in undifferentiated and synchronous activation of an overconnected functional network. Exogenous induction of synchronous activity with isoflurane anesthesia might dampen the FC difference between the two genotypes by increasing the FC in wild-type mice in a dose-dependent manner. This suggests that the FC abnormalities observed in awake Shank3b+/-mice might be masked or mitigated under anesthesia, highlighting the importance of considering brain state in the analysis of cortical network dynamics in autism models.

In summary, our study provides valuable insights into the complexity of autism-related cortical connectivity alterations from late development to adulthood. Our findings highlight behavioral anomalies at an early age (P20), a hyper-connected network involving barrel-field cortices during late adolescence (P45), and progressive deterioration over time until P90. We also demonstrated that brain state strongly influences cortical network dynamics: alterations are more pronounced in the awake state compared to under anesthesia, with divergent directionsobserved. We haveprovided unbiased and quantitativebiomarkers based on FC that hold high translational value for assessment, follow-up, and therapeutic development of PMS. These biomarkers will be fundamental for both preclinical and potentially clinical testing of novel therapeutic strategies.

## METHODS

### Ethical Statement

A total of 78 B6.129-Shank3^tm2Gfng^/J (3-12 weeks old) of both sexes were used. The Shank3B-knockout allele has a neo cassette replacing the PDZ domain (exons 13-16) of the Shank3 gene. Shank3B^+/-^ (n= 17) and Shank3B^+/+^ control littermates (n = 13) were used for mesoscopic calcium imaging (6 -12 weeks old). Shank3B^+/-^ (n= 38) and Shank3B^+/+^ control littermates (n = 11) were used for behavioral experiments (3 weeks old). All animals were housed in standard condition cages with a 12h light/dark cycle and food and water ad libitum. All experiment procedures were approved by the Italian Ministry of Health (aut n. 721/2020).

### Virus injection and intact-skull window

For CaMKII labeling with GCaMP7f, the viral construct ssAAV-PHPH.eB/2-mCaMKII*α*-jGCaMP7f-WPRE-bGHp(A) (1.3×10E13 vg/ml, volume: 50µL, Viral Vector Facility, CH) were diluted in 100µL of saline solution. A final volume of 150µL was intravenously injected in the retroorbital sinus of Shank3B^+/-^ and Shank3B^+/+^ mice under isoflurane anesthesia at P30, two weeks before imaging. One week before imaging the same animals were anesthetized with isoflurane (3% for induction, 1–2% for maintenance) and placed in a stereotaxic apparatus (KOPF, model 1900). Ophthalmic gel (Lacrilube) was applied to prevent eye drying; body temperature was maintained at 36°C using a heating pad and lidocaine 2% was used as local anesthetic. The skin and the periosteum were cleaned and removed, then bregma and lambda were signed with a black fine-tip pen. A custom-made aluminum head-fixing bar was placed behind lambda and the exposed skull was fixed using transparent dental cement (Super Bond C&B – Sun Medical). After the surgery, mice were recovered in a temperature- and humidity-controlled room, with food and water ad libitum.

### Wide-field microscopy setup

Wide-field imaging was performed using a custom-made microscope, as in Montagni and colleagues (Montagni et al. 2024). The microscope consisted of back-to-back 50 mm f/1.2 camera lenses (Nikon). To excite the GCaMP7f indicator, a 470 nm light source (LED, M470L3, Thorlabs, New Jersey, United State) filtered by a bandpass filter (482/18 Semrock, Rochester, New York, NY, USA) was deflected by a dichroic mirror (DC FF 495-DI02 Semrock, Rochester, New York, NY, USA) on the objective (TL2X-SAP 2X Super Apochromatic Microscope Objective, 0.1NA, 56.3 mm WD, Thorlabs). Reflectance images were acquired using a light source positioned at 45° incident to the brain surface (530 nm LED light, M530L4; Thorlabs, New Jersey, United State). Stroboscopic illumination (20Hz/LED) was used. The fluorescence and reflectance signals were selected by a bandpass filter (525/50 nm filter, Semrock, Rochester, New York, USA) and collected by a CMOS camera (ORCA-Flash4.0 V3 Digital CMOS camera / C13440-20CU, Hamamatsu). Images were acquired at 40Hz, with a resolution of 512×512 pixels (FOV of 11.5 × 11.5 mm).

### Habituation and awake imaging

After the post-surgical recovery period (3 days), mice were acclimatized to the head-fixation for two consecutive days (15 min a day/mouse) to gradually reduce anxiety and abrupt movements prior to data collection. 14 days after the injection, head-fixed imaging sessions were performed at different timepoints (P45, P60 and P90). Each imaging session consisted of 4 recordings (180 s-long) of spontaneous cortical activity in awake, resting-state mice that could freely move their limbs and weren’t engaged in any distinct tasks.

### Anesthetized imaging

The same mice that did the awake imaging were then anesthetized by isoflurane to investigate spontaneous cortical activity in two brain states classified according to the isoflurane concentrations in DEEP anesthesia (1.8 ± 0.1 %) and LIGHT anesthesia (1.1 ± 0.1 %). Each anesthesia level was maintained for 60 minutes, and recordings were consistently monitored to conserve a stable slow-oscillatory regime. Imaging sessions (180s-long, 5 repetitions) were performed at different timepoints (P45, P60 and P90). Deep and light anesthesia were recorded consecutively on the same imaging session per mouse, starting from the higher isoflurane concentration to the lower. During the whole anesthetic treatment, body temperature was maintained at 37°C by a feedback-controlled thermostatic heating pad.

### Image processing and data analysis

All data analyses were performed in MATLAB (MathWorks), Phyton, ImageJ, and Origin.

Image stacks for each animal collected from different sessions were registered using custom-made software, by considering bregma and λ position. An animal-specific field of view (FOV) template was used to manually adjust the imaging field daily. To dissect the contribution of each cortical area, we registered the cortex to the surface of the Allen Institute Mouse Brain Atlas (www.brain-map.org) projected to our plane of imaging. For each block, image stacks were processed to obtain the estimates of ΔF/F0. ΔF/F was computed for each pixel, where ΔF was the intensity value of that pixel in a specific time point and F was the mean fluorescence intensity of the signal across time. Then, hemodynamic correction was performed as described by Scott and colleagues (Scott et al. 2018). Briefly, using the ratiometric approach:

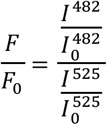

Where F/F0 is the final corrected GCaMP6 time series for a given pixel, I^482^ refers to the detected fluorescence signal, I^525^ is the reflectance signal.

In this study we focus on the analysis of the functional cortical network between Shank3B^+/-^ and Shank3B^+/+^. To analyze FC, a total of 22 ROIs were then selected (11 ROI for each hemisphere, 20×20 pixels). The abbreviations and extended names for each areas are as follows: MOs-a, anterior region of secondary motor cortex; MOs-p, posterior region of secondary motor cortex; MOp-a, anterior region of primary motor cortex; MOp-p, posterior region of primary motor cortex; SSp-bfd, primary somatosensory area, barrefield; SSp-tr, primary somatosensory area, trunk; SSp-fL, primary somatosensory area, forelimb; SSp-hl, primary somatosensory area, hindlimb; RSP, retrosplenial cortex; VISa, associative visual cortex; VISp, primary visual cortex. Throughout the text and figures, suffixes L and R were added to denote cortical areas of the left or right hemisphere, respectively (e.g., RSP_L_, RSP_R_). Correlation mapping was done for each subject by computing Pearson’s correlation coefficient between the average signals extracted from a ROI, with that of each other ROIs (time window: 180 s-long) and then averaged within a session. The averaged single-subject correlation maps were transformed using Fisher’s r-to-z transform and averaged across all animals within a group. Then, the group-related averaged maps previously obtained were re-transformed to correlation values (r-scores).

Differences between groups were calculated as r(Shank3B^-/+^)-r(Shank3B^+/+^) of the averaged correlation values, to visualize matrices of difference at all the time points. ROIs positions in FC graphs were arranged according to their anatomical coordinates. Lines and symbol sizes represented the level of correlated FC and the number of such connections associated with the ROI, respectively.

### Behavioral analysis

#### Open field

Open field test was conducted on postnatal day 20 (P20). Prior to the trial, mice were habituated to the experimental room for two hours. Then, mice were allowed to freely explore an empty arena (45 × 45 × 45 cm) for 10 minutes. The arena walls were white colored and smooth. Before the animal placement, the arena was washed with a 70% alcohol/water solution avoiding possible biasing effects from odor cues. The session was recorded with a camera(PointGrey flir Chameleon3, CM3-U3-13Y3C-CS), framerate was 30 fps. An experienced observer quantified the time spent self-grooming, which included behaviors such as cleaning the face, snout with paws, licking or biting the body. Videos were also automatically analyzed by Animal Tracker tool (ImageJ) to measure distance traveled and time spent in the center.

#### Movements during imaging session

At P45, to quantify movements during the imaging session, a camera (PointGrey flir Chameleon3, CM3-U3-13Y3C-CS) was orthogonally placed 10 cm in front of the mouse. The camera operated at a frame rate of 40 Hz, capturing images at a resolution of 512 by 512 pixels. This resolution provided sufficient detail to cover the frontal part of the mouse, allowing comprehensive analysis of movements. To well identify movements, visible illumination light at 630 nm was focused on the forepaws. Videos were then analyzed by an experienced observer to quantify time doing forepaw movements, including tapping and grooming.

### Brain slices preparation and electrophysiology recordings

Adult (p90) Shank3 mice were deeply anesthetized with isofluorane and decapitated for brain extraction. Acute coronal slices (350 μm thick) containing the primary somatosensory cortex (SSp-bfd) were cut with a vibroslaicer (Leica VT1000S, Leica Microsystem, Wetzlar, Germany) in ice-cold carboxygenated cutting solution containing in (mM): Sucrose (206), Glucose (25), NaHCO3 (25), KCl (2.5), NaH2PO4 (1.25), MgSO4 (3), CaCl2 (1). Slices recovered for a minimum of 1 hour in warm (32-34 °C), carbon-oxygenated, low-calcium artificial cerebrospinal fluid (aCSF) with the following concentrations (mM): NaCl (130), KCl (3.5), NaH2PO4 (1.25), NaHCO3 (25), Glucose (10), CaCl2 (1) and MgSO4 (2). Slices were individually transferred to a recording chamber placed under the objective of an upright microscope (Nikon Eclipse E600FN). During recordings, slices were continuously perfused with warm (32–34 °C) carbo-oxygenated, high-calcium aCSF solution composed of (in mM): NaCl (130), KCl (3.5), NaH2PO4 (1.25), NaHCO3 (25), glucose (10), CaCl2 (2) and MgSO4 (1). Whole -cell electrophysiological recordings were performed with borosilicate capillaries (Harvard Apparatus, London, UK) made by a vertical puller (Narishige PP830, Narishige International Ltd, London, UK) and, back -filled with K+ gluconate-based internal solution contained (in mM): 120 K-Gluconate, 15 KCl, 10 HEPES, 1 EGTA, 2 MgCl2, 5 Na2Phosphocreatine, 0.3 Na2GTP, and 4 MgATP (pH 7.3, 295–305 mOsm), resulting in a bath resistance of 3– 4 MΩ. Layer V Pyramidal neurons of SSp-bfd were studied by recording passive properties (membrane capacitance, membrane resistance, and resting membrane potential), intrinsic excitability (Current-voltage curves and action potential threshold), and spontaneous excitatory post-synaptic currents (sEPSCs) isolated by the application of 10 μM SR95531 (GABAA receptor blocker). Current-Action potential curves were obtained with the injection of increasing steps (500 ms) of depolarizing current ranging from -90 to +210 in 20pA increments. The threshold for action potential (AP) was determined at the base of the first spike. sEPSCs were recorded in voltage-clamp configuration holding the potential at -70 mV. All signals were sampled at 10 kHz and low-pass filtered at 2 kHz with an Axon Multiclamp 700B (Molecular Devices, Sunnyvale, CA, USA).

### Statistical analysis

For comparison between groups, One- or Two-Way ANOVA was used, followed by Bonferroni test. Group-level ROI-based FC differences between pre- and post-stroke groups were assessed by means of one-way repeated measure ANOVA followed by Tukey correction. Network Based Statistic (NBS) Toolbox in MATLAB was used to statistically assess functional network connectivity (Zalesky, Fornito, and Bullmore 2010; Nakai et al. 2023). We tested for both significantly higher and lower correlations. Differences were considered significant when p < 0.05. Errors are reported as Standard Error of Means, *p < 0.05, ** p < 0.01, *** p < 0.001. *Electrophysiological recordings* were analyzed using Clampfit 10.7 (Molecular Devices, Sunnyvale, CA, USA), Origin 2019 (OriginLab, Northampton, MA, USA) and GraphPad Software(San Diego, CA). All data were tested for normality distribution (D’Agostino-Pearson test, GraphPad Prism 7.0) before using parametric (Student’s t-test) or non-parametric (Mann-Whitney test) statistical test. *Behavioral test* Significant differences were tested with two-sample T-test.

## Acknowledgements

This work was funded by the Telethon Seed Grant for Spring 2022 (PHEM, GSA22E006), the Italian Ministry of Universities and Research PRIN 2022 (2022YCTLPL), and the THE Tuscany Health Ecosystem Project (ECS_00000017 MUR_ PNRR). Additional support was provided by Banca d’Italia and Fondo di Beneficenza di Intesa Sanpaolo through the RICONSIN Project (B/2021/0202), as well as the Fondazione Cassa di Risparmio di Firenze (CRF2020). Furthermore, this project benefited from the framework of Eurobioimaging (ESFRI research infrastructure) - Advanced Light Microscopy Italian Node.

## Contributions

ALAM conceived the study.

ALAM, LB, EM supervised the project.

ALAM, EM designed and interpreted experiment and computational analyses.

EM, MA, AMar, LC performed experiments, obtained tissue samples and generated data.

EM and LC conducted computational analyses.

EM, ALAM drafted the manuscript.

MA, LC, Amas, LB edited the manuscript.

ALAM, FSP, AMas, GM contributed funding and resources. All authors participated in the review of the manuscript.

